# Cortical isolation separates rhythmic synchrony from network integration in the human neocortex

**DOI:** 10.64898/2026.09.02.748777

**Authors:** Réka Bod, Kinga Tóth, Orsolya Farkas, Yossef Michaeli, Katalin Zsófia Tóth, Ágnes Kandrács, Katharina T. Hofer, Dániel Fabó, Boglárka Hajnal, Johanna Petra Szabó, Loránd Erőss, László Entz, István Ulbert, Lucia Wittner

**Author notes:** These authors contributed equally (István Ulbert, Lucia Wittner). Correspondence to: Lucia Wittner.

## Abstract

Human cortical activity reflects interactions between local recurrent circuits and distributed brain-wide inputs, but how these contributions shape cortical dynamics remains unclear. We compared oscillatory and single unit activity in laminar recordings from the same human cortical regions in eight patients across wakefulness, NREM sleep, and acute slices after surgical isolation. Gamma-band spike-field synchronization increased from wakefulness to sleep and isolated cortex, whereas population coupling, putative connectivity, network integration, and dynamical dimensionality declined. Laminar gamma sink-source organization persisted in isolation, indicating the presence of local gamma-generating mechanisms. Integration with histological data showed preserved tissue architecture, while neuronal density predicted population integration in vivo but not after isolation. Removing long-range inputs thus did not suppress cortical activity but shifted its organization toward stronger rhythmic coordination and reduced integration. Our findings suggest that neocortical microcircuits intrinsically generate coherent gamma activity, whereas network embedding supports the diverse, high-dimensional dynamics of the intact cortex.

## Introduction

The cerebral cortex operates through a continuous interplay between local recurrent circuitry and distributed interactions spanning cortical and subcortical networks. Dense excitatory and inhibitory connections within cortical microcircuits provide the substrate for neuronal firing and recurrent activity, while long-range inputs modulate these circuits across behavioral and physiological states (1–4). However, since local and distributed influences are inseparable in the intact brain, it remains difficult to determine which features of cortical dynamics are generated intrinsically by local circuitry and which depend on its embedding within the larger brain network.

Changes in brain state provide a physiological dimension along which this balance can be examined. Wakefulness is associated with relatively diverse and weakly correlated population activity (5–7), whereas NREM sleep increases neuronal synchrony and constrains population dynamics (8–11). These state-dependent changes indicate that cortical activity is not determined solely by the properties of individual neurons or local synaptic interactions, but also by the degree to which local circuits participate in distributed network activity (12). Nevertheless, physiological state changes do not remove long-range anatomical connectivity, leaving unresolved how much of this organization is retained when a cortical microcircuit is physically isolated.

Acute human cortical slices provide a complementary reduction of the intact system by preserving local synaptic circuitry while removing long-range afferent inputs, neuromodulatory influences, and behavioral context (13, 14). Human cortical slices can generate spontaneous population events and oscillatory activity, demonstrating that substantial network dynamics persist outside the intact brain (15–18). Whether these dynamics represent preserved physiological properties of human cortical microcircuits, remains still uncertain. In particular, it is unknown whether local circuits can retain coherent oscillatory coordination while losing the broader network integration characteristic of the intact cortex.

This question is important because synchronization and integration are not necessarily equivalent properties of a neural network (19, 20). Local recurrent excitatory-inhibitory interactions can generate temporally coordinated activity, whereas integration depends on interactions among distributed neuronal populations and may support a richer repertoire of population states (21–24). Consequently, removal of long-range inputs could increase local synchrony while simultaneously reducing the diversity and dimensionality of population activity. Whether such a dissociation occurs in human neocortex has not yet been directly tested.

The transition from wakefulness through NREM sleep to acute cortical isolation provides a particularly informative experimental axis. NREM sleep reduces large-scale cortical engagement while preserving anatomical connectivity, whereas surgical isolation removes long-range inputs altogether (25). Comparing these states within the same cortical regions therefore provides an opportunity to distinguish changes associated with reduced global drive from those produced by physical removal of distributed network interactions.

Here, we established a within-patient framework in which the same human cortical regions were studied during chronic intracranial monitoring and subsequently following therapeutic resection. Using identical 24-channel laminar microelectrodes, we recorded neuronal and local field activity during wakefulness and NREM sleep in the intact brain and from acute cortical slices prepared from the corresponding resected tissue. We combined single-unit electrophysiology, spike-field synchronization, population coupling, putative connectivity, network topology, population dimensionality, laminar current-source density, and quantitative histology to ask three related questions: (1) which electrophysiological features persist following cortical isolation, (2) how local synchronization and broader network integration are reorganized, and (3) whether these changes can be explained by alterations in local circuit structure or tissue integrity.

We found that along in vitro investigation neuronal single-unit activity proved to be comparable to the in vivo recording context, while strongly enhancing gamma-band spike-field synchronization. In contrast, population coupling, putative connectivity, network integration, and population dimensionality declined, despite preservation of laminar gamma sink-source organization and overall laminar architecture. As a consequence, cortical isolation revealed a dissociation between local rhythmic coordination and network integration: human cortical microcircuits retain the capacity to generate coherent oscillatory activity intrinsically, whereas their embedding within distributed brain networks supports the diverse and high-dimensional dynamics of the intact cortex. A minimal mechanistic model further identified external drive and local inhibitory architecture as complementary determinants of these two dimensions of cortical organization.

## Results

### Isolation alters intrinsic excitability while preserving baseline firing properties

To compare human cortical microcircuits in the intact brain with the same area-derived tissue obtained through surgical removal, we recorded laminar local field potentials (LFP) and single-unit activity using matched 24-channel microelectrode arrays during wakefulness, NREM sleep, and acute *in vitro* slice preparations from the same patients (n=8; Figure 1 a-c). We identified 546 well-isolated units across all conditions, classified into regular-spiking principal cells (RS-PCs), intrinsically bursting principal cells (IB-PCs), non-fast-spiking (NFS), and fast-spiking (FS) interneurons according to previously described methodology ((26), Figure 1d-e; Extended Data, Figure S2).

**Fig 1.**
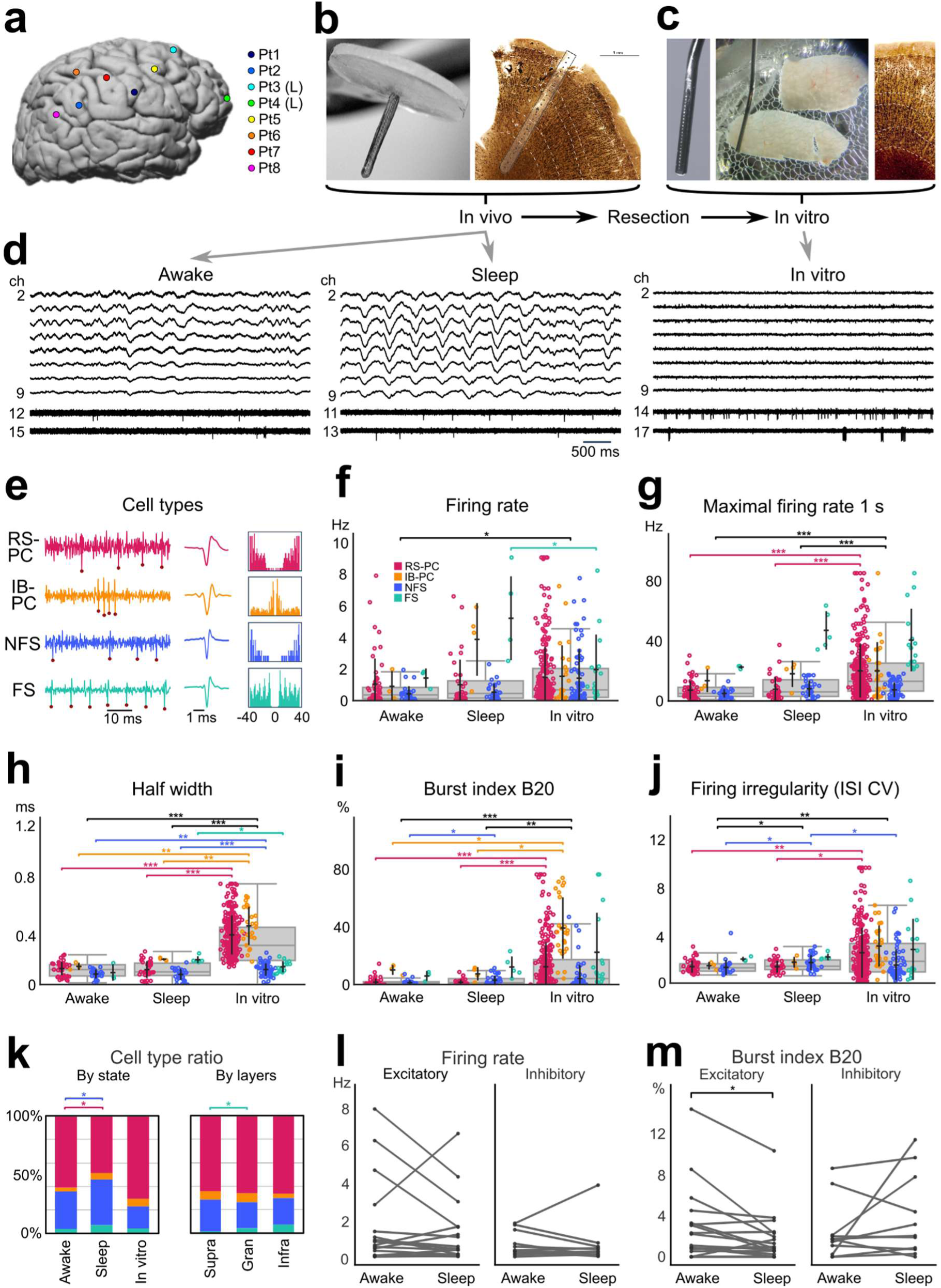
Experimental design and single-unit electrophysiological properties across wakefulness, NREM sleep, and acute cortical slices. (a-c) Experimental framework for matched recordings from the same cortical regions in the patients (n=8). (a) Representative MRI reconstruction showing the patient-specific location of the laminar microelectrode. (b) In vivo 24-channel laminar microelectrode and representative histological reconstruction of the electrode track across cortical layers, stained for NeuN. (c) Identical microelectrode used for acute in vitro recordings, representative cortical slice during recording, and corresponding NeuN staining. (d) Representative broadband local field potentials (LFPs, top 6 channels; 1-500 Hz) and high-pass-filtered multi-unit activity (bottom 2 channels, >500 Hz) recorded during wakefulness, NREM sleep, and acute in vitro conditions. (e) Representative single-unit classes identified from extracellular waveform morphology and firing properties: regular-spiking principal cells (RS-PCs), intrinsically bursting principal cells (IB-PCs), non-fast-spiking interneurons (NFS), and fast-spiking interneurons (FS). (f-j) Single-unit firing and waveform properties across recording conditions. (f) Mean firing rate. (g) Maximum firing rate within 1-s windows. (h) Action-potential principal halfwidth. (i) Burstiness index (B20). (j) Inter-spike-interval coefficient of variation (ISI CV). (k) Distribution of identified cell classes across recording conditions and layers; supra: supragranular layers (L1-3), gran: granular layer (L4), infra: infragranular layers (L5-6). (l-m) Longitudinal analysis of 42 units identified in both wakefulness and NREM sleep, showing changes in mean firing rate (l) and B20 (m). Group comparisons were performed with Mann-Whitney U tests and Benjamini–Hochberg correction. Data are presented as median/IQR boxplots with individual observations overlaid. *q<0.05, **q<0.01, ***q<0.001.

Isolation fundamentally altered intrinsic electrophysiological properties. Action potentials of both excitatory and inhibitory units substantially broadened *in vitro*, with median halfwidths increasing from ∼0.21 ms *in vivo* to 0.37 ms *in vitro* (Figure 1h; q<10⁻¹⁰ across all putative cell types). Despite these waveform changes, spontaneous baseline firing remained robust and differed only modestly across states. However, maximal firing rates (1-s maximal output) increased markedly after isolation, particularly among principal cells, reaching a median of 14 Hz *in vitro* compared to 6–7 Hz *in vivo* (Figure 1f-g; q<0.0001). Metrics reflecting the regularity of firing, including the inter-spike-interval coefficient of variation (ISI CV) and the proportion of ISIs shorter than 20 ms (burstiness index, B20), exhibited a progressive increase along the wakefulness-sleep-in vitro axis across all cell types. Firing irregularity of inhibitory NFS cells was higher in sleep compared to recordings obtained during wakefulness (Figure 1i-j) which came along with the different representation of cell types in the different conditions, the ratio of RS-PC/NFS being significantly lower in sleep compared to wakefulness. Furthermore, ratio of NFS cells increased towards deeper layers (Figure 1k). While tracking 42 distinct cells in both in vivo conditions, we did not find a monotonic firing rate tendency change in either putative excitatory or inhibitory classes, while burstiness index B_20_ in the same excitatory cells was significantly decreased in sleep (Figure 1 l-m).

### In vitro microcircuits exhibit paradoxical gamma hypersynchronization

Although isolation severed long-range inputs, it did not silence local rhythmic coordination, but on the contrary, it massively amplified it. While delta and theta phase-locking remained weak across all conditions examined (Extended Data, Figure S4A-B), gamma-band synchronization increased progressively along the wakefulness-sleep-in vitro axis. Low-gamma phase-locking values (PLV) rose from a median of 0.075 during wakefulness to 0.120 during NREM sleep, peaking at 0.369 in vitro (Fig. 2e). High-gamma PLV followed a similar trajectory (wake: 0.105, sleep: 0.144, *in vitro*: 0.414; Extended Data, Figure S4C). State-dependent tracking of identical units *in vivo* (n=42) confirmed that reduced arousal alone (NREM sleep) significantly enhanced both low- and high-gamma phase-locking relative to wakefulness, suggesting that isolation supports a physiological tendency toward gamma coordination already present during reduced cortical drive. Importantly, the hypersynchronization described previously preferentially recruited local inhibitory networks. *In vitro*, putative inhibitory neurons were five times as likely to exhibit significant gamma coupling, whereas excitatory populations showed a twofold increase. FS interneurons exhibited the strongest enhancement of gamma synchronization overall (all q<0.02), driving the highly synchronized *in vitro* state.

**Fig 2.**
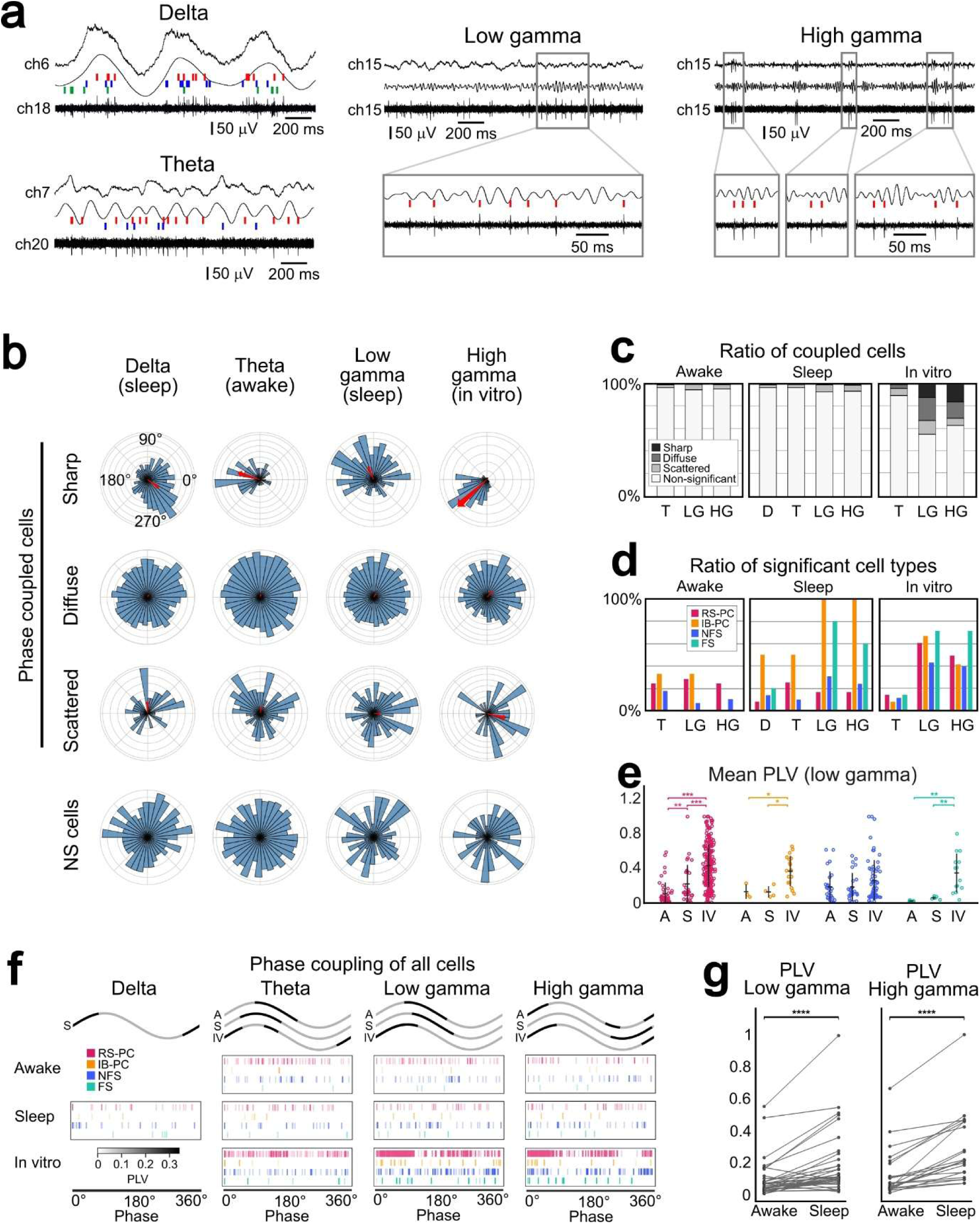
Spike–LFP phase locking across frequency bands and recording conditions. (a) Representative LFPs band-pass filtered in the delta (1–4 Hz), theta (4–8 Hz), low-gamma (30– 50 Hz), and high-gamma (50–80 Hz) bands with simultaneously recorded single-unit spikes. Insets show spike timing relative to individual oscillatory cycles. (b) Representative polar distributions of spike phases illustrating sharply coupled, diffuse, scattered, and non-significant phase-locking patterns (NS cells). Arrow length indicates phase-locking value (PLV) and arrow angle indicates preferred phase. (c) Proportion of single units assigned to each phase-locking pattern across recording conditions. (d) Proportion of units exhibiting significant spike–LFP phase locking in each frequency band. D: delta, T: theta, LG: low gamma, HG: high gamma. (e) PLV for low-gamma oscillations across recording conditions and putative cell classes. A: awake, S: sleep, IV: in vitro. (f) Preferred spike phases for significantly phase-locked units across frequency bands and recording conditions. Each raster represents one unit. Cell type is coded by color, coupling intensity is coded by color intensity (see explanation on the left side). Black line on the waves (upper panel) marks the preferred phase of the whole neuronal population examined (all cells pooled). A: awake, S: sleep, IV: in vitro. (g) Longitudinal comparison of low-gamma and high-gamma PLV in 42 units recorded during both wakefulness and NREM sleep. Statistical significance was assessed using Mann– Whitney U tests with Benjamini–Hochberg correction; paired data were analyzed using Wilcoxon’s test, and preferred firing phases relative to oscillatory cycles were evaluated with Rayleigh’s circular statistics. Data are shown as median ± IQR. *q < 0.05, **q < 0.01, **q < 0.001.

To determine if this enhanced gamma coupling is reflected in robust local current generation, we quantified laminar current-source density (CSD) and integrated sink strength (ISS). High-gamma sink-source organization remained remarkably stable across all states, with no significant differences in layer-specific ISS (Extended Data Figure S4D). Paradoxically, low-gamma sink strength was significantly reduced *in vitro* (Fig. 2f,g) despite the massive increase in single-unit phase-locking. This indicates that enhanced spike-LFP synchronization *in vitro* is uncoupled from bulk transmembrane current magnitude, likely reflecting a stereotyped, low-dimensional oscillatory regime rather than rich physiological network activity.

### Rhythmic coordination dissociates from structural network integration

Direct comparison of identical human cortical regions before and after surgical resection revealed a selective reorganization of cortical dynamics rather than a generalized loss of function. Population coupling, a proxy for functional network integration, declined nearly tenfold after isolation (median: awake=0.107, sleep=0.165, in vitro=0.039; q<0.0001) (Fig. 3b). This functional fragmentation was mirrored by a tenfold decrease in putative monosynaptic interactions, which dropped from 1.38% (15/1 088 tested pairs) in vivo to 0.18% (5/2 718 pairs) in vitro (Fig. 3a). PLV- and phase-based similarity networks (Fig. 3e) revealed that in vivo neurons showed a more interconnected population-coupling structure, whereas in vitro recordings exhibited stronger coherence in gamma-coupling profiles despite reduced population integration. Neurons in isolated cortex, this way, became more similar in their gamma-coupling behavior while becoming less embedded within the broader population activity. Across awake recordings, gamma-network closeness and degree were strongly anti-correlated (r = −0.94), indicating that gamma-synchronized hub cells are structurally peripheral (Fig 3d, Extendedd Data, Figure S4e). Population dimensionality metrics further supported this dissociation: eigenspectrum entropy of the covariance matrix, network participation ratio, and the fraction of total variance explained by the first principal component (PC1 variance) all showed a progressive reduction in participation ratio and eigenspectrum entropy, accompanied by an increase in PC1 variance from wakefulness to in vitro (Fig. 3d-f). These measures indicate a state-dependent reduction in dynamic dimensionality, in which rhythmic coordination persists despite structural disintegration.

**Fig 3.**
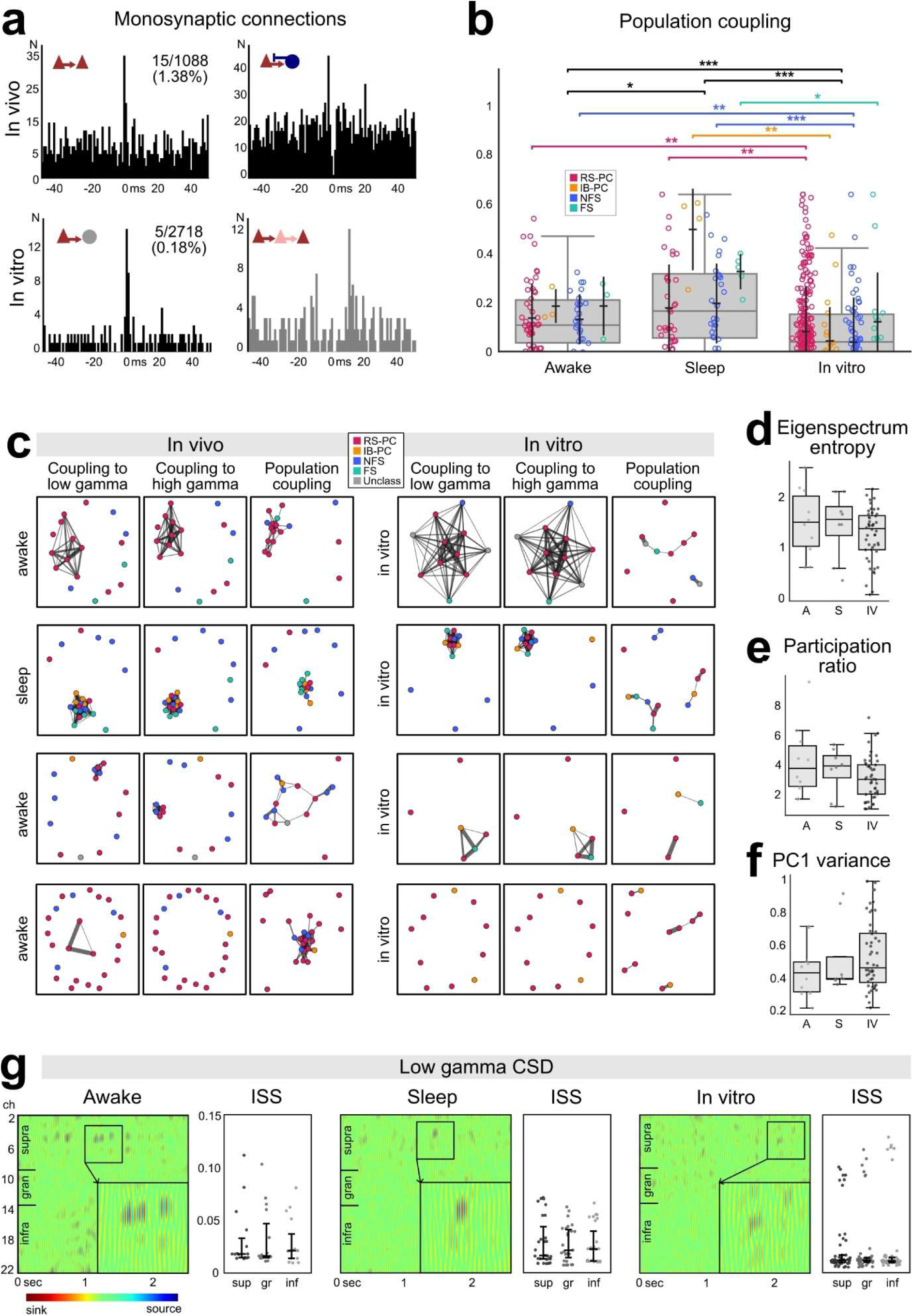
Functional network integration, population dimensionality, and laminar gamma activity across recording conditions. (a) Representative crosscorrelograms illustrating putative monosynaptic interactions and delayed population interactions between simultaneously recorded units. (b) Population coupling across wakefulness, NREM sleep, and acute in vitro conditions, shown separately for putative cell classes. (c) Representative functional network graphs constructed from low-gamma PLV, high-gamma PLV and population coupling. Note that neurons can be strongly (upper rows) or not coupled (bottom row) to gamma oscillation both in vivo (left) and in vitro (right) conditions. Population coupling in vivo involves most neurons of the microcircuit, which is dissociated into small networks in vitro. (d-f) Population covariance structure across recording conditions: eigenspectrum entropy (d), participation ratio (e), and variance explained by the first principal component (PC1; Ff. (g) Representative laminar low-gamma LFP and current-source density (CSD) profiles across recording conditions and corresponding integrated sink strength (ISS) across supragranular, granular, and infragranular layers. Group comparisons were performed using Mann-Whitney U tests with Benjamini-Hochberg correction. Data are displayed as median ± IQR boxplots with individual observations overlaid.*q < 0.05, **q < 0.01, ***q < 0.001.

### Structural preservation reveals a dissociation between architecture and functional integration

To determine whether the physiological reorganization observed after isolation could be attributed to tissue deterioration due to slice preparation and in vitro maintenance, we quantified neuronal and perisomatic inhibitory markers in the recorded neocortical samples using NeuN and parvalbumin (PV) immunohistochemistry, respectively (Fig. 4). NeuN-positive neuronal density exhibited the expected laminar organization, with substantially higher densities in supragranular than infragranular cortex (mean 1 241 versus 750 cells mm^-2^; q=7.14×10^-24^; Figure 4a). Importantly, overall neuronal density did not differ between tissue processed immediately after resection and tissue processed after completion of in vitro experiments (approx. 985 cells/mm^2^ in both groups; q=0.98), indicating the preservation of the overall neuronal population despite pronounced physiological reorganization.

**Fig. 4.**
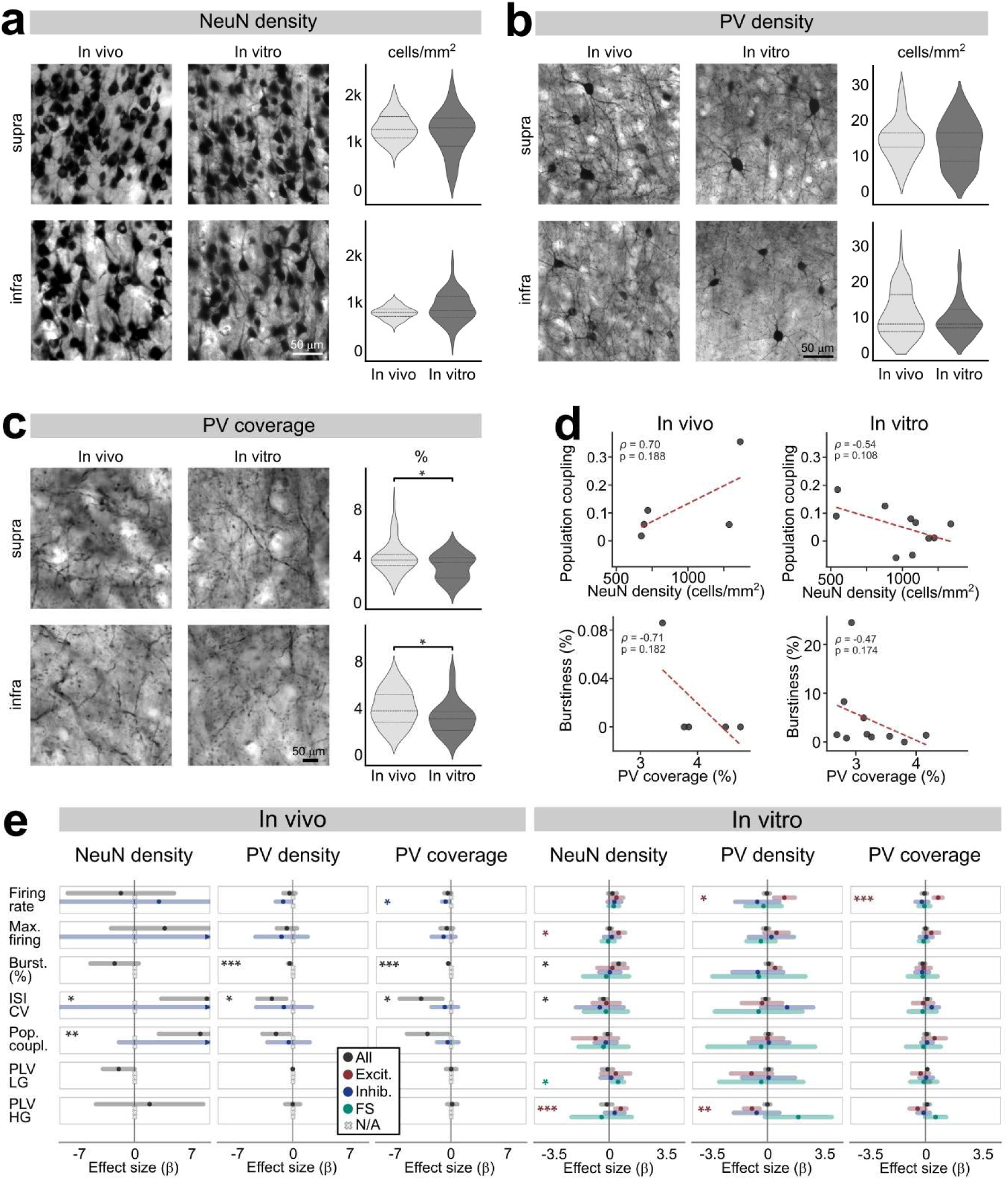
Histological organization and structure-function relationships before and after acute in vitro recording. (a-c) Representative immunohistochemical images and quantification of neuronal and inhibitory markers in supragranular and infragranular cortex before and after acute in vitro recording. (a) NeuN-positive neuronal density measured within 300 × 300 μm regions of interest, alongside with patientwise means overlaid on violin plots, in vivo (left panels) and in vitro (right panels), both in supragranular layers (L2-3, supra, upper panels), and infragranular layers (L5-6, infra, bottom panels). (b) PV-positive interneuron density measured within 500 × 500 μm regions of interest. (c) PV-positive process coverage within 500 × 500 μm regions of interest. (d) Spearman correlation relationships between histological measures and physiological variables, including NeuN density versus population coupling and PV process coverage versus burstiness. (e) Standardized coefficients from mixed-effects models testing associations between histological measures and single-unit physiological properties.Statistical testing employed Wilcoxon’s test, Spearman correlations and mixed-effects models: *p < 0.05, **p < 0.01, ***p < 0.001; n.s., not significant.

Markers associated with inhibitory circuitry exhibited more subtle changes. PV-positive interneuron density showed no significant layer-specific differences between post-in vivo and post-in vitro samples (Figure 4b; all q>0.05). In contrast, PV process coverage was modestly reduced after isolation in both supragranular and infragranular cortex (supragranular: 3.83% vs. 3.08%, q=0.040; infragranular: 4.30% vs. 3.45%, q=0.0017; Figure 5C), suggesting partial remodeling of inhibitory neuropil despite preserved overall cellular architecture.

**Fig 5.**
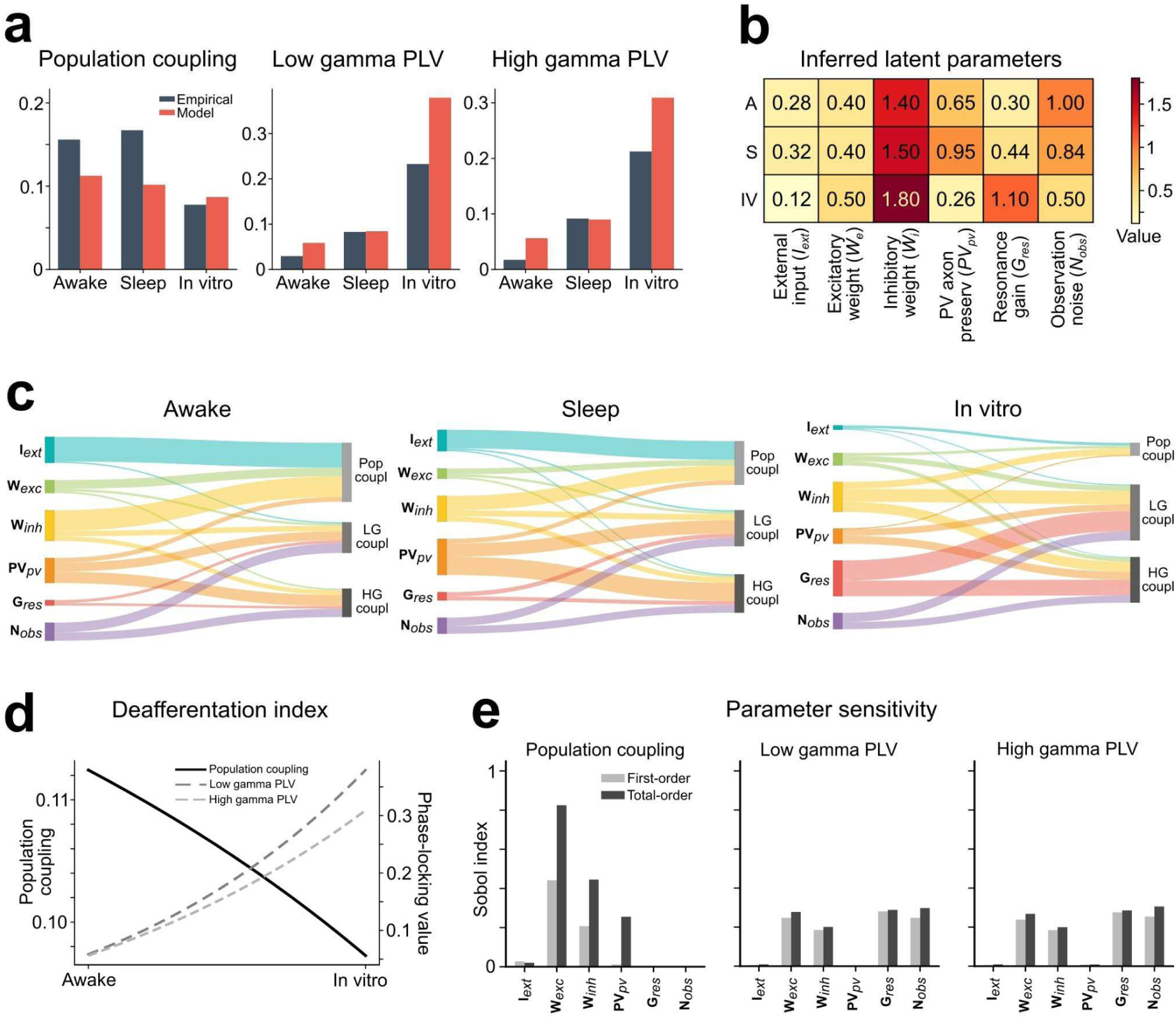
Minimal mechanistic model of the dissociation between population coupling and gamma phase locking. (a) Empirical and model-derived population coupling, low-gamma PLV, and high-gamma PLV across awake, NREM sleep, and acute in vitro conditions. (b) Optimized values of the model parameters across recording conditions. (c) Contribution of model parameters to population coupling and gamma phase locking based on the model decomposition. Abbreviations are explained on subfigure (b), Pop coupl: population coupling; LG and HG coupl: low gamma and high gamma coupling, respectively. (d) Continuous model trajectory obtained by interpolation between the optimized awake and in vitro parameter sets. (e) First-order Sobol sensitivity indices quantifying the contribution of individual model parameters to population coupling, low- and high-gamma phase locking.

We next asked whether local cytoarchitecture predicted microcircuit function. In tissue recorded in vivo, neuronal density emerged as a strong positive predictor of population coupling in mixed-effects analyses (standardized β=7.76, q=8.18×10^-253^; Figure 5E). Regions containing higher neuronal densities therefore exhibited stronger integration into ongoing population activity. PV-related measures were similarly associated with reduced excitatory firing and burst generation, consistent with inhibitory regulation of local network excitability (Figure 4f, g).

Strikingly, these relationships were markedly altered following isolation. Whereas neuronal density positively predicted population coupling in the intact brain, this association was substantially weakened and reversed in isolated tissue (Figure 4e). Correspondingly, rank correlations between NeuN density and population coupling shifted from positive in vivo (ρ=0.70) to negative in vitro (ρ =-0.54), although these correlations did not individually reach significance owing to the limited number of histological samples. Similar weakening of structure–function relationships was observed for PV-related measures and burst dynamics (Figure 4f).

Moreover, multivariate analyses also indicated that these associations extended to firing and oscillatory properties of the local neuronal circuits. In vivo, PV density and coverage were negatively associated with ISI variability across the whole population, consistent with a greater inhibitory organization associated with more regular firing (Fig. 4e). In vitro, higher NeuN density was associated with stronger high-gamma PLV and greater maximal firing output (over 10 s) of putative excitatory neurons, whereas PV density had an inverse association with excitatory-cell high-gamma PLV. PV density and coverage were, interestingly, positively associated with excitatory firing measures, possibly suggesting that a preserved inhibitory architecture supports elevated firing output, even without further increasing gamma-rhythmic coupling.

These findings indicate that the transition from the intact brain to isolated cortex occurs without substantial neuronal loss. More than merely reflecting degeneration, isolation selectively disrupts the coupling between local cytoarchitecture and functional network organization. Thus, neuronal density predicts integrative dynamics only when cortical microcircuits remain embedded within distributed brain networks.

### A minimal mesoscopic model identifies complementary mechanisms of the dissociation

To determine whether this dissociation can arise from a small set of biophysically interpretable parameters, we constructed a minimal analytical mesoscopic model (STAR Methods). The model managed to recover the central paradox: population coupling declined monotonically from the awake to the in-vitro state, matching empirical values within 28 % (awake) and 12 % (in vitro), while both low- and high-gamma PLV increased along the same axis (Fig. 5a). Optimized parameters revealed a coherent progression: loss of external drive, reduction of PV axonal preservation, and compensatory rise in local resonance gain (Fig. 5b). Sankey decomposition of wakefulness, slow-wave sleep and in vitro showed that population coupling was dominated by the external-drive term, whereas gamma PLV was controlled by resonance gain and PV axonal survival (Fig. 5c). Continuous interpolation between awake and in-vitro parameter sets produced a smooth decline in population coupling accompanied by a monotonic rise in both gamma bands (Fig. 5d). Sobol sensitivity analysis confirmed that external drive is the principal determinant of population coupling, while resonance gain and PV axonal survival dominate gamma phase-locking (Fig. 5e). Thus the two axes of the paradox are controlled by dissociable circuit parameters.

## Discussion

Direct comparison of identical human cortical regions before and after surgical resection revealed a selective reorganization of cortical dynamics rather than a generalized loss of function. Isolation preserved spontaneous firing and laminar gamma generators, while increasing single unit action potential-gamma synchronization, but simultaneously reduced population coupling, putative connectivity, network integration, and population dimensionality. NREM sleep occupied an intermediate position between wakefulness and isolated cortex across several measures. These findings collectively indicate that cortical synchrony and network integration are related but dissociable dimensions of cortical organization.

This dissociation is particularly evident in the simultaneous increase in gamma phase locking and decrease in population coupling. Oscillatory synchrony is often considered a mechanism for coordinating neuronal communication and may facilitate the selective routing of information between neuronal populations (27, 28). However, phase locking to a local oscillation and participation in population-wide activity quantify different aspects of network organization. A neuron may become more precisely aligned to a common gamma phase while becoming less correlated with the broader population (29, 30). In the present data, this distinction was explicit: after isolation, neurons showed stronger gamma phase locking despite weaker population coupling, fewer putative interactions, reduced graph integration, and lower dimensionality. As a result, the increased synchrony did not indicate preserved or enhanced network integration, but more convincingly, it suggested that neuronal activity became increasingly organized around a restricted local temporal reference while losing access to the diverse activity patterns generated through distributed network interactions.

This interpretation is also consistent with the preservation of laminar gamma generators after isolation. Gamma activity is strongly associated with recurrent interactions between excitatory neurons and fast-spiking, parvalbumin-positive (PV+) interneurons, whose rapid inhibition provides a powerful mechanism for temporal coordination (31–33). Spike timing relative to oscillatory phase can provide a precise temporal organization of neuronal output, even when the broader network does not support extensive interaction diversity (34). The present findings extend this framework to human neocortex by showing that local gamma coordination can become stronger when the cortical microcircuit is removed from the intact brain.

Our histological observations make this finding particularly informative. PV-positive process coverage was reduced after acute slice preparation, consistent with partial truncation of the extensive dendritic and axonal arborizations of PV+ interneurons during tissue isolation. Such structural disruption would intuitively be expected to weaken PV-mediated temporal coordination and thereby reduce gamma synchronization (35, 36). Instead, gamma phase locking increased markedly, particularly among fast-spiking interneurons. This apparent paradox indicates that the presence of an intact PV network is not, by itself, sufficient to predict the magnitude of gamma synchronization measured at the level of spike-field coupling. More plausibly, local recurrent circuitry may retain sufficient inhibitory architecture to generate rhythmic coordination even after partial structural compromise, while the removal of competing long-range inputs may increase the relative dominance of the remaining local oscillatory mechanism. PV+ interneurons are particularly well positioned to support this process because of their efficient perisomatic inhibitory functions, their extensive reciprocal connectivity, fast membrane dynamics, and strong recruitment by excitatory inputs (33, 37–40).

The reduction in population coupling may therefore also reflect, at least in part, the combined consequences of long-range deafferentation and local inhibitory reorganization. Population coupling measures the extent to which individual neurons participate in broader population fluctuations, and therefore depends on the network context in which local activity is embedded ((41–43). Removal of corticocortical, thalamocortical, and neuromodulatory inputs directly eliminates sources of distributed covariation (44). Concurrent disruption of PV processes may further alter the spatial and temporal spread of local activity, potentially fragmenting interactions that previously linked neuronal populations across cortical layers (45, 46). The present data cannot establish a causal contribution of PV process loss to the reduction in population coupling, but the parallel changes in PV-positive process coverage, population coupling, and gamma phase locking suggest that local inhibitory remodeling should be considered alongside loss of long-range input when interpreting the isolated state.

Our electrophysiology-histology associations further implicated local inhibitory architecture in the control of both spike output and temporal coordination through oscillations. In vivo, greater PV density and coverage were associated with more regular population firing, whereas in the isolated cortex, greater PV density was associated with increased excitatory firing output but reduced excitatory-cell gamma phase locking. These associations do not establish direct causality, particularly given the limited number of histological samples, but suggest that the relationship between PV-mediated inhibitory architecture, spike output and rhythmic coordination depends strongly on network context.

The distinction between oscillatory and population coupling also clarifies why strong synchronization can coexist with reduced interaction structure. A network dominated by a common oscillatory drive can produce highly coordinated spike timing without maintaining the heterogeneous pairwise interactions characteristic of the intact cortex (47, 48). In such a regime, many neurons become aligned to the same temporal structure, increasing phase synchrony while reducing the number of independently varying dimensions of population activity. The observed reduction in participation ratio, eigenspectrum entropy, and PC1-independent variance is consistent with this interpretation. Isolation, in this context, appears to shift cortical dynamics from a distributed, high-dimensional regime toward a locally resonant, low-dimensional regime.

The progression from wakefulness through NREM sleep to isolated cortex provides further support for this framework (49). Sleep preserved anatomical connectivity but altered the effective balance of large-scale interactions, and several measures (including gamma synchronization and population dimensionality) shifted toward the isolated state. Complete isolation amplified these changes. This progression suggests that the in vitro state should not be interpreted simply as a pathological or non-physiological preparation, but as an experimentally accessible endpoint of reduced network embedding. Local circuits become increasingly dominant as external drive and distributed interactions are progressively diminished.

The findings may also have broader relevance to disorders in which local circuit integrity and large-scale integration become uncoupled. Alterations of PV+ interneurons and gamma synchronization have been implicated in epilepsy, schizophrenia, and several neurodegenerative disorders, although the mechanisms and direction of these changes differ substantially between diseases (50–52). The present experiments provide a useful mechanistic distinction: altered gamma synchronization does not necessarily imply preserved network integration. More generally, pathological or degenerative processes that disrupt long-range connectivity, inhibitory architecture, or both could produce complex combinations of preserved local rhythmicity and impaired large-scale communication. This possibility should be tested directly rather than inferred from oscillatory measures alone.

Finally, the minimal model supports the interpretation that these observations arise from interacting mechanisms rather than a single physiological change. Reducing external drive reproduced the progressive loss of population coupling and dimensionality, whereas changes in local inhibitory circuitry and resonance parameters influenced gamma phase locking. In the light of these, the empirical phenotype is most parsimoniously understood as the combined consequence of loss of distributed afferent drive and reorganization of local inhibitory circuitry. However, it should be noted that the model does not establish that PV-process loss is the causal source of the enhanced gamma coupling, but rather, it demonstrates how local resonance can increase while population integration declines when these mechanisms are partially decoupled.

Taken together, these findings demonstrate that the capacity of human cortical microcircuits to generate coherent oscillatory activity is remarkably robust to loss of large-scale network inputs. What is lost with isolation is not neuronal coordination per se, but the capacity to embed that coordination within a diverse set of distributed interactions. This way, the intact cortex appears to operate through a balance between local resonance and large-scale integration: recurrent microcircuits provide the machinery for temporal coordination, whereas distributed inputs expand the variety of states in which neuronal circuits can function.

## Methods

### Patients and Ethics

Eight patients with pharmacoresistant focal epilepsy underwent invasive presurgical monitoring followed by therapeutic cortical resection at the National Institute of Clinical Neurosciences (now Clinic for Neurosurgery and Neurointervention, Semmelweis University), Budapest, Hungary. All patients contributed both chronic intracranial recordings and subsequent acute cortical slice recordings. Patient demographic and clinical characteristics, including age, sex, pathology, seizure onset zone, and recording and resection locations, are summarized in Table S1. Written informed consent was obtained from all participants in accordance with the Declaration of Helsinki. All procedures were approved by the Hungarian Medical Scientific Council and the relevant institutional ethical committees (20680-2/2011/EKU).

### Intracranial Electrode Implantation and In Vivo Recordings

Twenty-four channel linear microelectrodes (150 μm distance between the contacts) were implanted into the neocortex of eight pharmacoresistant epileptic patients, below the clinical subdural grid electrodes (53). Recordings were part of the clinical investigation aiming to identify the seizure focus and the eloquent areas prior to surgical therapy, and were performed as described earlier (26). Briefly, continuous video-EEG investigation was made for 5–7 days. Electrocorticogram (EcoG) was recorded from the clinical grid (20–48 channels, mastoid reference) was recorded concurrently with the patient video using the standard system of the hospital. The laminar microelectrode was implanted perpendicular to the cortical surface, and the local field potential gradient (LFPg) was simultaneously recorded in all layers of the neocortex with EcoG. Electrode locations were verified using clinical MRI and intraoperative photographic documentation. Microelectrode contacts 1-12 were usually located in the supragranular layers (L1-3), contacts 13-16 in the granular layer (L4) and contacts 17-24 in the infragranular layers (L5-6). The exact intracortical location of the microelectrode was reconstructed with post hoc anatomy in three patients, whereas contact locations were extrapolated in the remaining five patients based on the above reconstructions and the in vitro recordings (see below).

In all eight cases the analyzed recordings originated from cortical regions that were subsequently resected and used for post hoc in vitro recordings. This enabled the comparison of activity recorded during presurgical monitoring with spontaneous neuronal activity recorded from corresponding acute cortical tissue.

Only physiologically stable interictal recordings were included. For each patient, approximately 30-min artifact-free periods of resting wakefulness and NREM sleep were selected for each condition. No seizures occurred within 1 hour before or after the analyzed recording periods. Wakefulness and NREM sleep were identified using the spectral characteristics of the LFP recordings together with the contemporaneous clinical and electrophysiological assessment documented by the treating epileptology team. Clinically recorded ECoG signals were not included in the analyses presented here.

### Tissue Resection and Acute Slice Preparation

Immediately after therapeutic resection, cortical tissue was transferred to the laboratory within the same building and maintained continuously in oxygenated solution. For tissue preparation, specimens were initially immersed in ice-cold oxygenated sucrose solution containing (in mM): 248 sucrose, 26 NaHCO₃, 1 KCl, 1 CaCl₂, 10 MgCl₂, and 10 D-glucose, continuously bubbled with 95% O₂ and 5% CO₂. 500-µm-thick neocortical slices were prepared perpendicular to the cortical gray-matter axis with a Leica VT1000S vibratome (RRID:SCR_016495). Slices were allowed to recover for approximately 1 h at 32–34°C before recording.

Recordings were performed in an interface chamber maintained at 32–35°C with continuous perfusion of oxygenated artificial cerebrospinal fluid (ACSF; 95% O₂/5% CO₂). ACSF contained (in mM): 124 NaCl, 26 NaHCO₃, 3.5 KCl, 1 MgCl₂, 1 CaCl₂, and 10 D-glucose.

### In Vitro Electrophysiological Recordings

The same 24-contact laminar geometry was used for acute slice recordings, with the probe positioned perpendicular to the cortical surface along the L1-6 axis (26). The laminar microelectrode was adapted to the in vitro setup by changing the flexible cable to a long rigid stainless steel tube from the electrode contacts to the preamplifier. Slices were systematically mapped at approximately 300-400 µm intervals to identify recording locations with detectable spontaneous cellular activity and adequate laminar coverage. Only recordings obtained under physiological ACSF, without pharmacological manipulation were included in this study.

Signals were acquired using the matched laminar probe and the same acquisition system with the same parameters as for the chronic intracranial recordings (0.01-10 kHz band-pass filtering; 20-kHz sampling rate, 16-bit digitization). The probe was positioned perpendicular to the pia mater reaching all cortical layers, and recordings were retained when spontaneous cellular activity was clearly detectable and remained physiologically stable throughout the analyzed epoch.

For each patient, acute slice recordings were obtained from perielectrode-excised tissue corresponding to the cortical region sampled during presurgical monitoring. This experimental design provided a within-patient comparison between activity recorded in the intact brain and spontaneous activity emerging in the corresponding isolated cortical tissue.

### Spike Sorting and Unit Classification

Action potentials were detected and initially sorted using Kilosort4 (54) and manually curated using Phy (https://github.com/cortex-lab/phy). Clusters were subsequently refined using the custom interactive SpikeWise tool (https://github.com/rekabod/spikewise), which enabled inspection and curation of cluster waveform morphology, autocorrelograms, refractory-period structure, and firing statistics. Single-unit isolation required inter-spike interval (ISI) refractory period violations (< 1.5 ms) to remain below 0.1% of total spikes. Clusters were further curated by removing individual events that were judged not to belong to the corresponding unit based on waveform morphology and temporal structure.

Putative principal cells and interneurons were separated using extracellular action-potential waveform kinetics. Spike half-width was the primary discriminating feature, with repolarization slope, trough-to-peak latency, waveform symmetry, and peak-to-trough amplitude ratio used as additional features. Principal cells were classified as regular-spiking principal cells (RS-PCs) or intrinsically bursting principal cells (IB-PCs) based on autocorrelogram structure, burst propensity, waveform characteristics, and manual inspection. Interneurons were subdivided into fast-spiking (FS) and non-fast-spiking (NFS) populations using a two-component Gaussian mixture model fitted to maximum firing rate, with an intersection threshold of 20.1 Hz. Final classifications were manually verified. Units were confirmed as identical across wakefulness and NREM sleep if they were detected on the same or neighouring physical channel (stabile due to the silicone sheet on the top of the in vivo electrode) and maintained consistent waveform dynamics. Waveform stability was quantified by calculating Pearson correlation coefficients (r > 0.9) on mean 2 ms spike templates across states, alongside cross-state stability in peak-to-trough amplitude, half-width duration, and principal component feature space clustering. Representative waveform measurements, classification features, and calibration procedures are provided in Figure S2 in the Extended Data.

### Spike Waveform and Firing Statistics

Action potentials were extracted from the assigned recording channel and characterized using waveform and firing features. Waveforms were aligned to the negative voltage trough. Action-potential half-width was calculated as the full width at half-maximum of the most prominent waveform: HW_i_ = *t*_post,50%_ − *t*_pre,50%_. Repolarization slope was calculated as the maximum positive derivative of the waveform during the return from the inverse peak toward baseline:

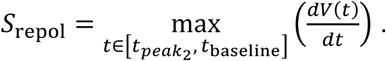

For unit *i*, mean firing rate was calculated as *R̅_i_* = *N_i_*/*T*, where *N_i_* is the number of detected spikes and *T* is the recording duration in seconds. Maximum firing rates were calculated as 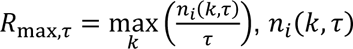 is the number of spikes emitted by unit *i* within the *k*-th window of duration *τ*, with *τ* =1 or 10 s.

Temporal firing structure was quantified from consecutive inter-spike intervals (ISIs), defined as *I_k_* = *t_k_*_+1_ − *t_k_*, where *t* denotes the timestamp of the *k*-th spike. ISI coefficient of variation was calculated as 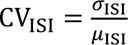. Burstiness was quantified using the *B20* index, defined as the proportion of consecutive ISIs shorter or equals 20 ms, defined formally *B*20 = 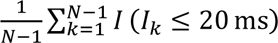, where *I*(·) is the indicator function. Thus, higher *B20* values indicate a greater fraction of high-frequency inter-spike intervals. Burstiness index B% was defined as sequence of three action potentials occurring within 20 ms was classified as a potential burst:

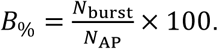

Bursts containing more than three action potentials could extend beyond 20 ms provided that each consecutive group of three action potentials occurred within a 20-ms window. The first and last action potentials of a burst had to be preceded and followed, respectively, by a 20-ms silent period, according to (26). Units were classified according to waveform morphology and firing dynamics as RS-PCs, IB-PCs, NFS interneurons, and FS interneurons. Full definitions of waveform and firing measures are provided in the Extended Data.

### Spectral Analysis and Epoch Detection

LFPg signals were analyzed to characterize oscillatory activity across recording conditions. Spectral power was estimated using Welch’s method with 4-s windows and 50% overlap. Oscillatory epochs were identified within predefined frequency bands: delta (0.5–3.5 Hz), theta (4–8 Hz), low-gamma (30–50 Hz), and high-gamma (50–80 Hz).

Band-specific activity thresholds were determined using paired reference recordings and receiver operating characteristic (ROC) analysis with Youden’s J statistic. When ROC-based separation was insufficient, percentile-based thresholds were applied using the consensus distribution across recording channels. Only epochs exceeding the predefined spectral threshold were included in subsequent phase-locking analyses. For further details, consult Extended Data and Figure S3.

Wakefulness and NREM sleep epochs were selected from physiologically stable recordings based on LFP spectral characteristics and the clinical electrophysiological assessment. In vitro recordings were analyzed during stable spontaneous activity under physiological ACSF.

### Spike–LFP Phase Locking

Spike–LFP phase locking was quantified separately for low-gamma (30–50 Hz) and high-gamma (50–80 Hz) activity. LFP phase was obtained from the analytic signal of band-pass-filtered LFPg using the Hilbert transform: *Z*_LFP_(*t*) = *A*(*t*)*e^iϕ^*^(*t*)^, where *ϕ*(*t*) is the instantaneous phase.

For each unit, phase-locking value (PLV) was calculated from the phases sampled at spike times: 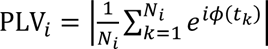

The preferred phase was calculated from the circular mean of spike-associated phases:

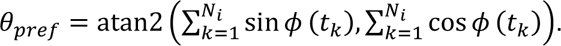

Statistical significance was assessed using both the Rayleigh test and a circular-shift permutation procedure. Epochs were retained for analysis only when they contained at least 10 spikes and satisfied the predefined significance criteria.

### Population Coupling and Functional Connectivity

Population coupling quantified the relationship between the activity of an individual unit and the activity of the surrounding neuronal population. Spike trains were binned at 1-s resolution, and for each unit the Pearson correlation was calculated between its firing rate and the population rate after excluding that unit (Okun et al., 2015): *c_i_* = corr(*r_i_*(*t*), *P*_–*i*_(*t*)), where *P_−i_*(*t*) = Σ_*j≠i*_*r_j_*(*t*).

Pairwise functional relationships were additionally assessed from cross-correlograms of spike trains. Putative monosynaptic interactions were identified from short-latency peaks or troughs in the 1-5 ms range relative to a 20-50 ms baseline, using a threshold of ≥4 standard deviations above baseline. These interactions were classified as putative excitatory or inhibitory according to the polarity of the correlogram feature.

### Network Construction and Graph Analysis

Functional networks were constructed separately for each recording session from pairwise neuronal activity relationships. Non-negative pairwise weights were retained above a predefined threshold to form weighted undirected graphs *G* = (*V*, *E*, *W*), where nodes represented neuronal units and edges represented functional relationships. The specific mathematical formulations are provided in Extended Data, Table S5.

### Population Dimensionality and Signal Complexity

Population activity dimensionality was quantified from the covariance matrix of neuronal firing activity. Principal component analysis (PCA) was used to obtain the eigenvalue spectrum *λ*_1_ ≥ *λ*_2_ ≥ ⋯ ≥ *λ_n_* ≥ 0. Three complementary measures were calculated. The participation ratio (PR) quantified effective population dimensionality: 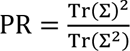., where Tr(Σ) is the sum of the eigenvalues. The proportion of variance explained by the first principal component (PC1) quantified the dominance of the largest collective mode: 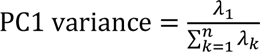. Eigenspectrum entropy quantified the distribution of variance across dimensions: *H*_λ_ = 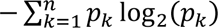, where 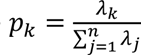 signifies probability distribution. PR and eigenspectrum entropy were interpreted as complementary measures of population dimensionality, whereas PC1 variance quantified concentration of activity into the dominant collective mode.

### Laminar Current-Source Density and Integrated Sink Strength

Laminar current-source density (CSD) was estimated from the second spatial derivative of the local field potential (LFP) across the regularly spaced recording contacts. CSD profiles were aligned to the cortical laminar organization and analyzed separately across supragranular, granular, and infragranular recording depths:

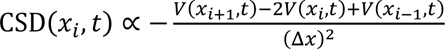, where the numerator is the second spatial derivative of the voltage and (Δ*x*)^2^ normalizes the derivative by the square of the electrode spacing. As we recorded the LFPg, CSD was calculated as the first spatial derivative using standard techniques (53). To quantify the magnitude of transmembrane current generation, integrated sink strength (ISS) was calculated by integrating the magnitude of negative CSD components (CSD_–_(*x*, *t*)) within predefined laminar and temporal windows: ISS*_L_* = 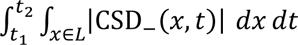. Layer-specific ISS was used to compare the strength and distribution of current sinks across wakefulness, NREM sleep, and acute slice recordings.

### Histological Processing and Quantification

Tissue blocks obtained immediately after resection and neocortical slices collected after acute electrophysiological recording were fixed and processed for immunohistochemistry as described previously (55). Sections were cut at 60 µm. Neuronal somata were labeled with mouse monoclonal anti-NeuN (1:2,000, EMD Millipore, RRID: AB_2298772), and parvalbumin-positive interneurons with mouse monoclonal anti-PV (1:7,000, Swant, RRID: AB_10000343). Immunoreactivity was visualized using established immunohistochemical procedures (17). Images were acquired using either a Nikon N-STORM System or a Leica DM 2500 (RRID:SCR_020224). Quantification was performed using the (https://github.com/rekabod/neu_roi_ranger) image-analysis workflow. The analysis included thresholding and cell annotation modules, with manual correction available for obvious segmentation errors.

ROIs were assigned independently by two experienced biologists/histologists after inspection of the complete tissue section, without initially examining the tissue at high magnification. Approximately five ROIs per cortical layer were placed in supragranular and infragranular cortex for each tissue condition, with ROI locations adjusted to provide representative tissue sampling. Image brightness and contrast were adjusted during image preparation to obtain comparable visualization of tissue and staining across sections.

NeuN- and PV-positive somata were quantified within the predefined ROIs and converted to densities: 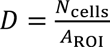, converted to cells/mm^2^. PV process coverage was defined as the proportion of each ROI occupied by PV-immunoreactive cellular elements: PV coverage = 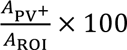. PV-positive area was determined by standardized thresholding, with limited manual addition or removal of detected elements when necessary. The image-analysis parameters were applied consistently within each analysis set. ROIs from the same patient, tissue condition, and cortical layer were averaged before statistical testing, resulting in the patient-level or tissue-level aggregate, rather than individual ROIs, was treated as the experimental unit. For details, see Figure S6 in Extended Data.

#### Minimal analytical mesoscopic model

In our efforts to examine mechanisms that could dissociate population coupling from gamma phase locking, we developed a minimal mesoscopic network model with 100 nodes representing neurons distributed along a normalized cortical axis. Node positions were defined 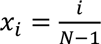, *where i* = 0, …, *N* − 1. The spatial arrangement reproduced approximate cortical laminar proportions, with 35% of nodes assigned to supragranular, 20% to granular, and 45% to infragranular positions. Nodes were assigned as 80% excitatory and 20% inhibitory.

Connectivity was distance dependent and followed an exponential decay function, *P_ij_* = 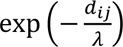, where *d_ij_* = |*x_i_* − *x_j_*|, is the distance between nodes and λ is the axonal footprint length constant. Recurrent connections were separated into local (*d*_ij_ ≤ 0.35) and long-range (*d*_ij_ > 0.35) components: *W*_local_ = *W*_raw_ ⊙ *M*_local_, *W*_long_ = *W*_raw_ ⊙ *M*_long_; *W*_raw_ is the distance-dependent weight matrix and *M*_local_ and *M*_long_ are binary distance masks. Excitatory and inhibitory recurrent gains were represented by *g_E_* ∈ [0.5, 3], *g_I_* ∈ [0.5, 3.5] and *λ* ∈ [0.05,0.6] respectively. Baseline values of λ were approximately 0.45 for wakefulness, 0.30 for NREM sleep, and 0.08 for the in vitro condition, representing progressive restriction of long-range connectivity following cortical isolation. Network activity was modeled as a noise-driven multivariate Ornstein-Uhlenbeck process. The stationary covariance matrix Σ was obtained by solving the continuous Lyapunov equation, *A*Σ + Σ*A^T^* + *Q* = 0; where *A* is the effective network dynamics matrix and *Q* is the noise covariance matrix. External drive was modeled as sparse input to supragranular nodes. Population coupling was calculated as the mean absolute correlation between each node and the population activity excluding that node: *C*_pop_ = 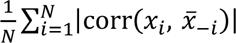, where *x_i_* is the activity of node and *x̅_-i_* is the mean activity of all other nodes. Local gamma synchrony was represented by an explicit resonance component whose amplitude depended on the local excitation-inhibition balance, PV axonal survival, and external-drive suppression. Low-gamma (30–50 Hz) and high-gamma (50–80 Hz) phase-locking values were generated by saturating the resonance amplitude with an observation-noise term. PV axonal survival was represented by a parameter *s_PV_* ∈ [0, 1], where lower values progressively reduced PV-mediated recurrent connectivity to mimic axonal truncation following acute cortical isolation.

The six latent parameters were external drive (*g_ext_*), recurrent excitation (*g_E_*), recurrent inhibition (*g_I_*), PV axonal survival (*s_PV_*), resonance gain (*g_res_*) and observation noise (*σ_obs_*): ***θ*** = (*g_ext_*, *g_E_*, *g_I_*, *s_PV_*, *g_res_*, *σ*_obs_).

Parameters were independently optimized for awake, NREM sleep, and in vitro conditions using differential evolution against three primary empirical targets: population coupling, low-gamma PLV, and high-gamma PLV. Participation ratio and PC1 variance were used as secondary validation measures. A continuous deafferentation trajectory was generated by linear interpolation between the optimized parameter sets, and first-order Sobol sensitivity indices were calculated to quantify the relative contribution of each parameter to population coupling and low-gamma PLV. For a model output *Y*, the first-order Sobol index for parameter was calculated as 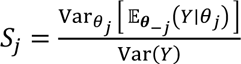, where ***θ****_-j_* denotes all parameters other than *θ_j_*. Parameter contributions to population coupling and gamma phase locking were visualized using a variance-based Sankey decomposition. Global Sobol sensitivity analysis calculated the total-effect index (*ST_i_*) for each minimal model parameter. Relative variance contributions were scaled proportionally to represent total direct and interactive parameter effects. Flows originate from individual biophysical parameters (e.g., synaptic decay constants, maximum conductances), pass through intermediate population dynamic nodes, and terminate at primary targets (population coupling strength and gamma phase-locking value).

### Statistical analysis

All statistical analyses were performed in Python 3.11 using NumPy, SciPy, statsmodels, and relevant analysis-specific packages. Code to replicate the main results and figures of the manuscript is available at https://github.com/rekabod/invivo_invitro_paper_code. Statistical tests were selected according to the experimental design and distribution of the data. Paired continuous measurements were compared using Wilcoxon signed-rank tests when appropriate, and unpaired comparisons using Mann-Whitney U tests. Categorical proportions were compared using the χ² test (as appropriate). For comparisons involving more than two experimental conditions, pairwise post hoc comparisons were performed using the corresponding paired or unpaired tests, with multiple comparisons controlled using the Benjamini-Hochberg false-discovery-rate (FDR) procedure. Adjusted p values are reported as q values, with statistical significance defined as q<0.05.

In analyses containing repeated measurements from multiple units or ROIs within patients, hierarchical structure was accounted for using linear mixed-effects models, with patient included as a random effect and experimental condition and relevant biological factors included as fixed effects. Where appropriate, recording session or tissue condition was included as an additional random effect. Standardized regression coefficients (β) are reported together with 95% confidence intervals and FDR-adjusted q values.

Associations between histological and physiological measurements were assessed using Spearman’s rank correlation. Where multiple measurements were obtained from individual patients, mixed-effects models were used to avoid treating repeated measurements as independent observations. For network and population analyses, the recording session or patient-level aggregate was used as the relevant experimental unit according to the analysis.

Circular statistics were performed using the Rayleigh test together with circular-shift permutation testing for spike-LFP phase locking, when the first test showed significant differences. Phase-locking analyses were restricted to epochs containing at least 10 spikes and satisfying the predefined significance criteria. Exact sample sizes, including the number of patients, recording sessions, units, unit pairs, and histological ROIs contributing to each analysis, are reported in the corresponding figure legends and Extended Data, Excel Spreadsheet. All analysis code is available in the project repository.

## Acknowledgements

The authors thank Domokos Meszéna for his valuable comments on the study design and for insightful suggestions regarding statistical analysis and data interpretation.

## Funding

This research was funded by the National Research, Development, and Innovation Office, grant nos. PD143380 (to Á. K.), FK129120 (to K. T.), K137886, Advanced 150799 (to L.W.); by the Hungarian Brain Research Program, grant no. NAP2022-I-2/2022 (to I.U.); by the János Bolyai Research Scholarship (to K. T.) of the Hungarian Academy of Sciences, and by the EU: FLAG-ERA Joint Transnational Call 2021, VIPattract grant (to L. W.), EIC Transition 2024, FlairVision grant (to W.L.). The scientific research and results published here were reached with the Doctoral Student Scholarship Program of the Co-operative Doctoral Program of the Ministry of Innovation and Technology (to R. B.), and the Semmelweis 250+ Graduate Student Fellowship of the Semmelweis University (to R. B.).

## Data availability

Given that patients at the onset of the study did not consent to public release of their data, raw data will be made available from the corresponding author (L.W.) upon request to protect patient privacy and consent.

## Author contributions (CRediT)

Conceptualization, R.B., L.W., I.U.; Methodology, L.W., R.B., I.U.; Investigation, R.B., O.F., Y.M., K.Zs.T., K.T., Á.K., K.H., D.F., J.Sz., B.H., L.E., L.Er., L.W.; Formal analysis, R.B.; Software, R.B.; Data curation, R.B., O.F., Y.M. K.Zs.T.; Resources, L.E., L.Er., D.F., I.U., L.W.; Visualization, L.W., R.B.; Supervision, I.U., L.W.; Writing -original draft, R.B., L.W; Writing -review & editing, R.B., L.W.

## Declaration of Conflict of Interest

The authors declare no conflict of interest.

## Declaration of generative AI and AI-assisted technologies in the writing process

During the preparation of this work, the authors used the OpenAI ChatGPT 5.6 model to improve clarity and enhance the overall language quality of the manuscript. A locally hosted distribution of Qwen 3.6 was employed for code development and statistical analysis, supplemented by the International Brain Laboratory (IBL AI Agent, https://github.com/int-brain-lab/ibl-ai-agent) project for refining coding structure and elaborating mesoscopic model principles. After using these tools, the authors thoroughly reviewed and edited all content and take full responsibility for the integrity and accuracy of the published article.

## Notes

### Competing Interest Statement

The authors have declared no competing interest.

