## Supplementary Data for "Cortical isolation separates rhythmic synchrony from network integration in the human neocortex"

### Extended Data for „Cortical isolation separates rhythmic synchrony from network integration in the human neocortex”

### Figures, Tables and Legends

**Table S1. Clinical demographic and surgical targeting data, Related to STAR Methods.** Summary of patient-specific characteristics for the cohort (n = 8) from which identical human cortical microcircuits were tracked across in vivo wakefulness, NREM sleep, and in vitro acute slice preparations. "Investigated cortical region" denotes the exact anatomical location of the 24-channel laminar microelectrode array during both physiological and isolated states.

| Pt ID | Sex | Age | Epilepsy onset (yrs) | Diagnosis | Investigated cortical region | Suspected seizure onset zone |
|---|---|---|---|---|---|---|
| Pt1 | M | 18 | 5 | Focal cortical dysplasia | Right postcentral gyrus (parietal) | right postcentral gyrus |
| Pt2 | F | 31 | 18 | Focal cortical dysplasia IIb with balloon cells | Right postcentral sulcus (parietal) | right parieto-temporal region |
| Pt3 | M | 48 | 39 | Focal cortical dysplasia IIb with tuberous sclerosis | Left frontocentral | left fronto-medial gyrus |
| Pt4 | F | 46 | 35 | Focal cortical dysplasia Ib/IIa | Left superior frontal gyrus | fronto-temporal alternating sites |
| Pt5 | F | 18 | 15 | Focal cortical dysplasia IIb with balloon cells | Right frontal | right supplementary motor area |
| Pt6 | M | 30 | 27 | Focal cortical dysplasia IIb with balloon cells | Right parietal | right supramarginal gyrus |
| Pt7 | M | 32 | 23 | Focal cortical dysplasia IIb with balloon cells | Right parietal | right frontal inferior gyrus |
| Pt8 | M | 42 | N/A | Tuberous sclerosis | Right parietal | right supramarginal gyrus |

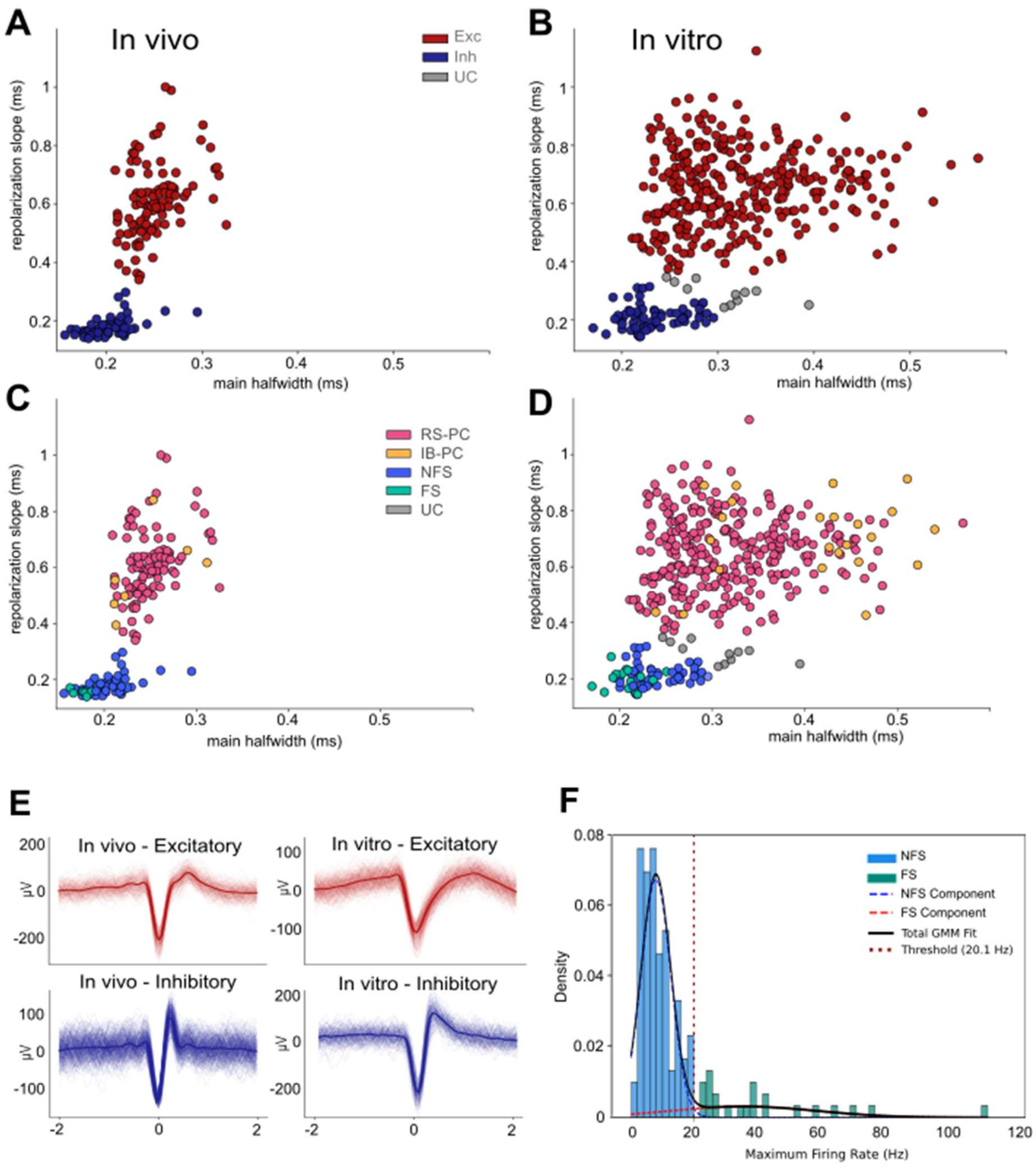


**Figure S2. Electrophysiological celltype classification and state-dependent waveform dynamics, Related to STAR Methods and Figure 1e (Results).**

(A, B) Scatter plots of single-unit extracellular action potential parameters: main halfwidth (x-axis) versus repolarization slope (y-axis) for units recorded in vivo (A) and in vitro (B), color-coded by broad functional class: putative excitatory neurons (red) and putative inhibitory neurons (blue).

(C, D) Fine-grained cell-type segmentation across in vivo (C) and in vitro (D) conditions, separating recorded units into regularly spiking principal cells (RS-PC, magenta), intrinsically bursting principal cells (IB-PC, orange), non-fast-spiking interneurons (NFS, sky blue), and fast-spiking interneurons (FS, teal).

(E) Overlay of representative single-unit extracellular waveform averages across in vivo and in vitro states, highlighting the physiological broadening of action potentials following cortical isolation.

(F) Separation of interneuron subpopulations using Gaussian Mixture Modeling (GMM) on 1s-windowed maximal firing rates, establishing an empirical decision boundary at 20.1 Hz to isolate fast-spiking (FS) units.

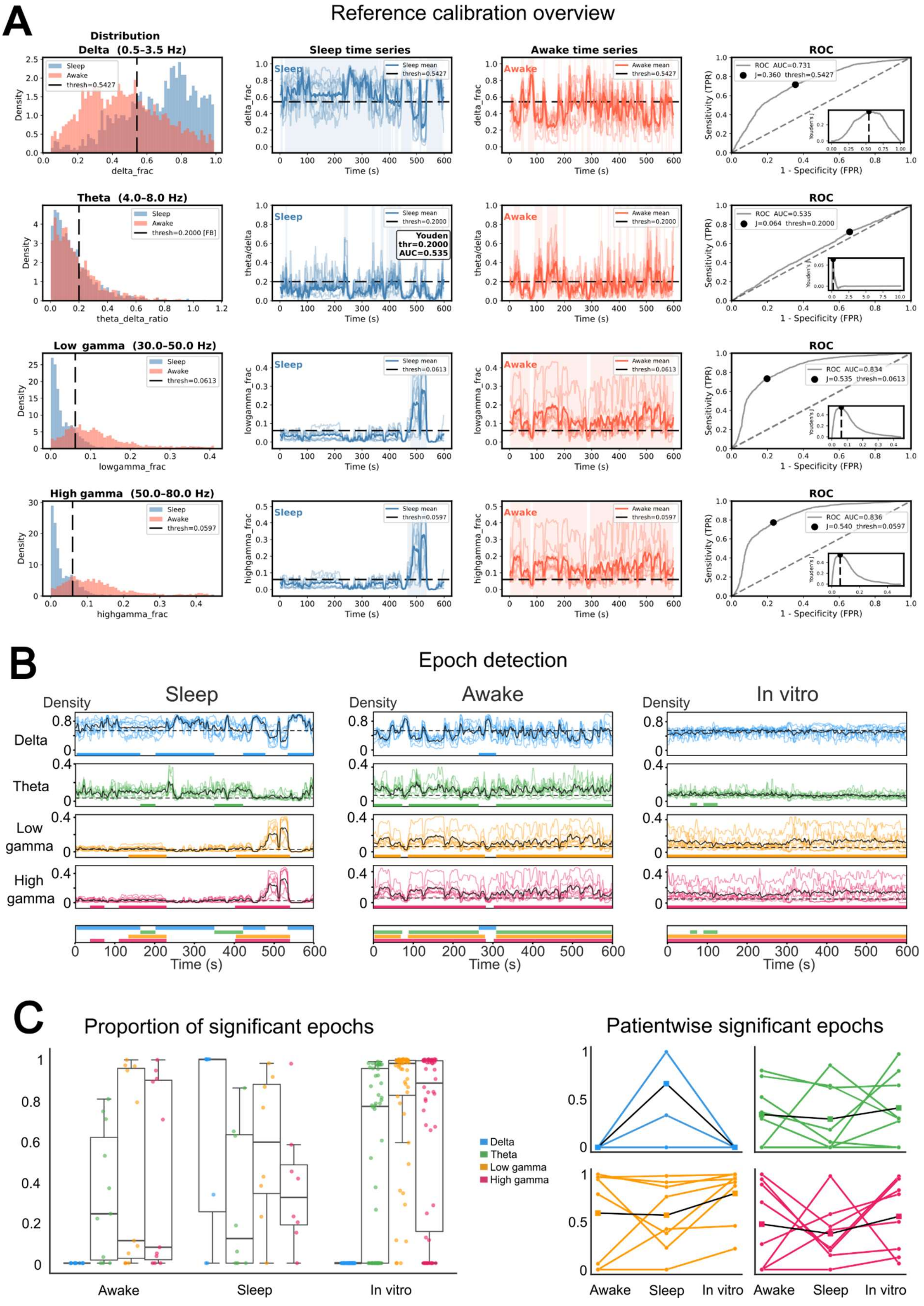


**Figure S3. Automated Threshold Calibration and Quantification of Oscillatory LFP Epochs Across Physiological and Experimental States, Related to Figure 2 (Results).**

(A) Schematic of spectral feature extraction and subject-matched threshold calibration. Left column: Empirical metric probability density distributions for paired NREM sleep (orange) and quiet wakefulness (slate blue) baseline reference states in a representative subject. Dotted vertical lines denote the optimal classification threshold calculated by maximizing Youden's J statistic. Middle panels: Representative time-frequency power spectral density heatmaps computed across paired reference recordings during NREM sleep (2nd column) and quiet wakefulness (3rd column) used to derive fractional band power distributions and band power ratios. Right column: Receiver operating characteristic (ROC) curves evaluating binary state-classifier performance for candidate spectral metrics. Sensitivity is plotted against 1-Specificity. Shaded circular markers indicate optimal cutoff points. For candidate metrics exhibiting weak discrimination; horizontal dashed threshold line), the detection engine automatically executes a 95th percentile fallback routine on reference baseline distributions to establish robust detection boundaries.

(B) Representative 10-minute continuous LFP traces demonstrating automated epoch detection across experimental states (wakefulness, NREM sleep, and in vitro slice preparation). Faint traces depict individual recorded microelectrode channels; thick black lines represent the across-channel mean LFP signal. Subject-calibrated spectral thresholds are indicated by dashed horizontal lines. Periods where spectral power continuously exceeds threshold for longer than minimum duration criteria are highlighted (shaded regions), marking significant oscillatory epoch detection.

(C) Quantification of significant LFP band epoch proportions within single recording sessions. Boxplots depict single-recording fractional ratios of total recording duration spent in significant oscillatory states across canonical frequency bands ($\delta$: 0.5-4 Hz, $\theta$: 4-8 Hz, $\gamma_{low}$: 30-50 Hz, and $\gamma_{high}$: 50-80 Hz) across conditions. Boxes indicate median and interquartile range (IQR); whiskers denote 1.5×IQR; individual circles represent single recording sessions. (D) Patient-level cohort aggregation of significant LFP epoch ratios. Line plots connecting individual patient averages across states display the state-dependent evolution of significant oscillatory coverage (n=8 patients). Note that δ oscillations were only measured during NREM sleep and the elevated γ-band oscillatory persistence in isolated in vitro cortical microcircuits.

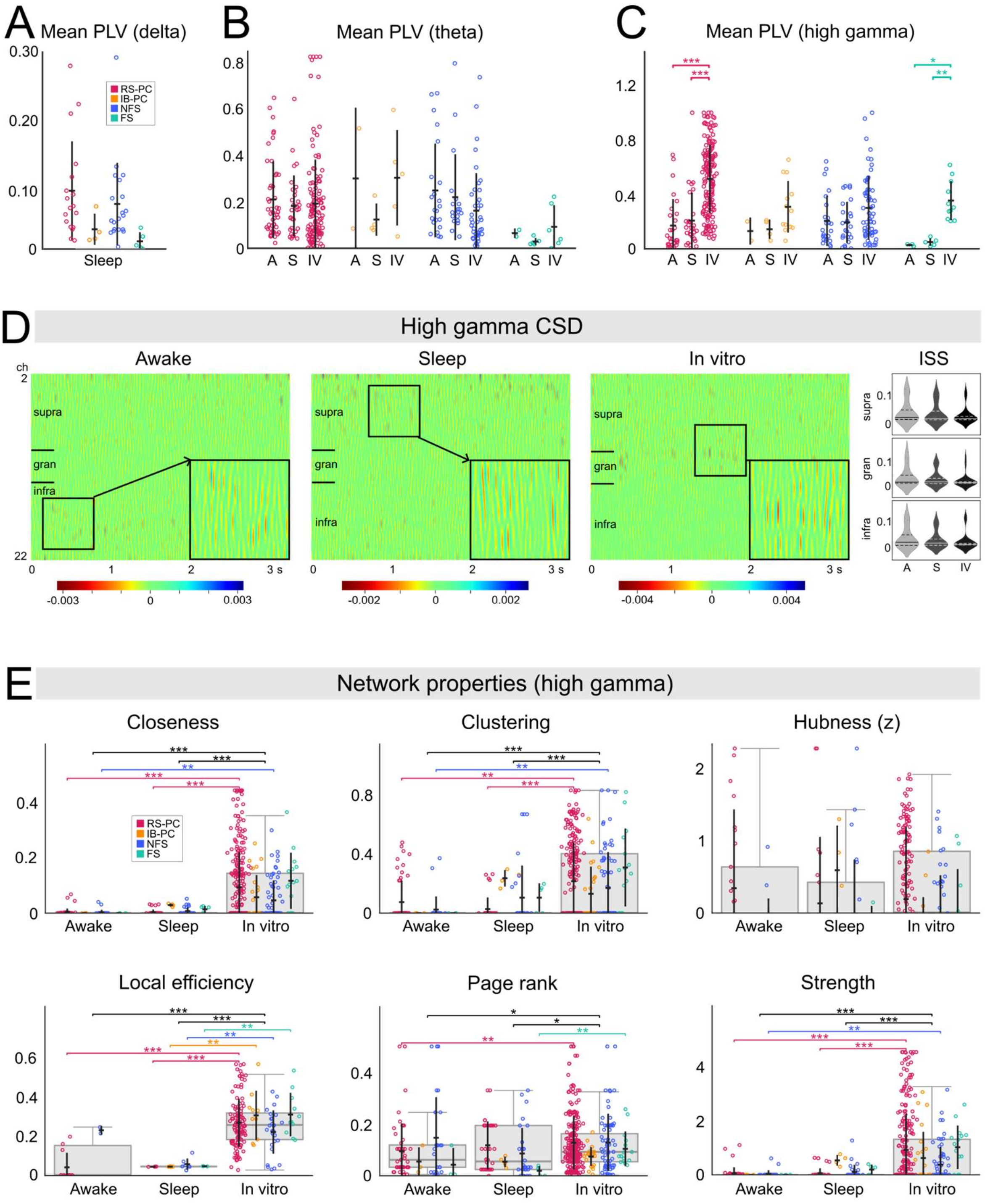


**Figure S4. State-dependent oscillatory phase-locking, preserved high-gamma current generators, and functional network topology, Related to Figure 3 (Results).**

(A) Mean Phase-Locking Values (PLV) in the δ-band across functionally defined cell types (RS-PC, IB-PC, NFS, FS) strictly during NREM sleep. (

B) Mean θ-band PLV across wakefulness, NREM sleep, and in vitro slice conditions.

(C) Mean high-γ (50-80 Hz) PLV across all three physiological and isolated states. For panels A–C, individual single-unit values are overlaid as scatter points on a horizontal median line with whiskers indicating the interquartile range (IQR).

(D) Representative high-γ Current Source Density (CSD) spatial maps extracted from recording snippets across wakefulness, NREM sleep, and in vitro states. Accompanying violin plots quantify the Integrated Sink Strength (ISS) across supragranular (supra), granular (gran), and infragranular (infra) layers, demonstrating that the spatial profiles of high-γ generators remain preserved regardless of global state or isolation.

(E) Single-unit functional network properties across conditions and cell types. Boxplots overlaid with individual cell-level scatter points display closeness centrality, clustering coefficient, standardized hubness (Z-score), local efficiency, PageRank centrality, and node strength. Color-coded horizontal significance bars and asterisks denote significant differences between state and cell-type groups. Data distributions are represented as medians ± IQR. Statistical evaluations utilize Mann-Whitney U with Benjamini-Hochberg False Discovery Rate (FDR) correction. Significance brackets reflect FDR-corrected q-values (*q < 0.05, q < 0.01, ***q < 0.001).

**Table S5, Graph Metrics Mathematical Formulations, related to Figure S4 E.**

| Metric | Mathematical formulation | Biological / Structural Interpretation |
|---|---|---|
| Closeness centrality | $CC_i = \frac{n-1}{\sum_{j \neq i} d\,(i,j)}$ | Proximity of unit $i$ to all network nodes, based on shortest path distance $d(i,j)$. |
| Clustering coefficient (weighted) | $C_i = \frac{1}{S_i(k_i - 1)} \sum_{j,h} \frac{w_{ij} + w_{ih}}{2} a_{ij} a_{ih} a_{jh}$ | Local interconnectivity and triad tightness surrounding unit $i$. |
| Hubness (z) | $H_i = \frac{S_i - \bar{S}}{\sigma_S} + \frac{CC_i - \overline{CC}}{\sigma_{CC}}$ | Composite score identifying driver units exceeding population variance in strength and closeness. |
| Local efficiency | $E_{loc,i} = \frac{1}{k_i(k_i - 1)} \sum_{j,h \in G_i, j \neq h} \frac{1}{d_{G_i}(j,h)}$ | Fault tolerance and functional communication efficiency in the neigh-borhood $G_i$ of unit $i$ if unit $i$ is removed. |
| PageRank | $PR_i = \frac{1-d}{N} + d \sum_{j \in M(i)} \frac{PR_j}{L(j)}$ | Relative global influence based on links to other high-degree nodes (damping factor = 0.85). |
| Node strength | $S_i = \sum_{j \in V} w_{ij}$ | Total functional coupling weight of unit $i$ across the population. |

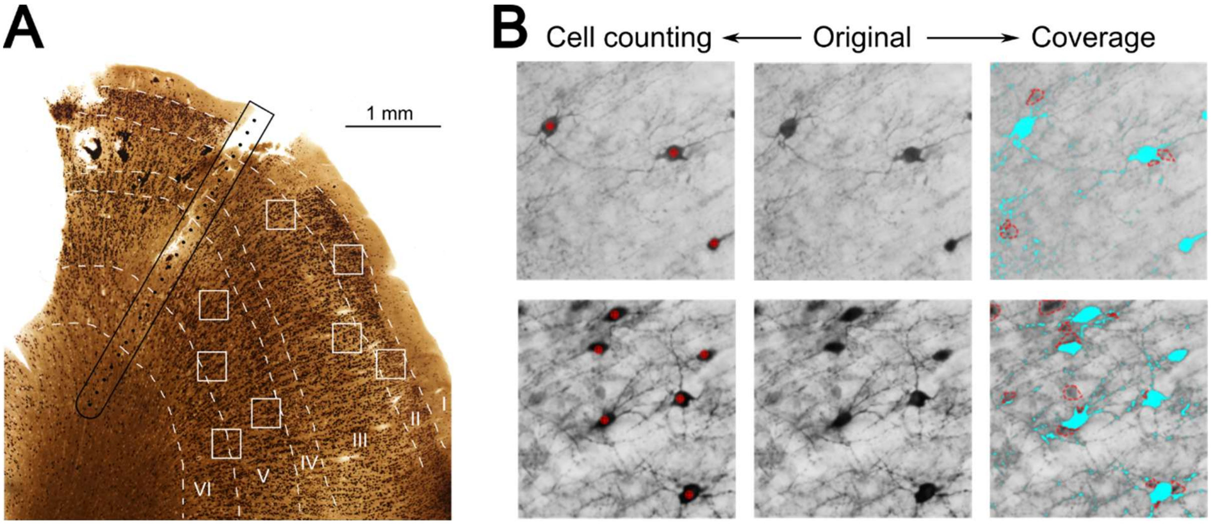


**Figure S5. Histological track reconstruction and semi-automated region of interest (ROI) quantification framework, Related to STAR Methods**.

(A) Representative histological reconstruction of the resected neocortical slice. The physical laminar microelectrode track is rendered visible in black, alongside the superimposed Regions of Interest (ROIs) framed as white, utilized for layer-specific cellular density counting across the supragranular and infragranular compartments. Dashed lines signify reconstructed anatomical layers of the neocortex, ranging layers I-VI.

(B) The custom graphical user interface (GUI) of the ROI annotator utilized for the quantitative analysis. The framework displays 500 × 500 $\mu m^2$ field-of-view samples of parvalbumin (PV) immunostaining. The interface integrates a direct cell counter module, the original raw micrograph, and a spatial coverage module. To ensure high-fidelity quantification across variable tissue samples, the coverage module allows for expert-supervised adjustments of exposure, contrast, and dynamic bimodal mask thresholding. Light blue represents PV coverage, whereas red dashed lines show areas excluded due to out of focus.
